# Drought escape as an adaptive strategy across an aridity gradient in wild sunflower

**DOI:** 10.64898/2026.05.26.728009

**Authors:** Rajesh Neupane, Brennan R. Silva, Louis S. Santiago, Kate Ostevik

## Abstract

**Premise:** Drought is intensifying in many regions under climate change, affecting plant growth, phenology, and resource allocation. Plants cope with water limitation through strategies such as escape, avoidance, and tolerance, but how these strategies are reflected in trait variation across natural drought gradients remains poorly understood. We examined how drought-related traits vary across a drought gradient in wild sunflower (*Helianthus annuus* L.).

**Methods:** We conducted a common-garden experiment using 19 wild sunflower populations collected across Southern California spanning a climatic moisture deficit gradient. We quantified growth, phenology, morphology, biomass allocation, and physiological traits and evaluated their relationships with climatic moisture deficit and soil variables using regression and correlation analyses.

**Results:** Populations from drier environments exhibited coordinated shifts in life-history strategy, combining faster growth, earlier flowering, and increased reproductive output. These changes were coupled with reduced allocation to roots and structural tissues and diminished investment in traits that buffer plant water status, as reflected in declines in relative water content and higher (less negative) leaf water potentials. Ordinal modeling further indicated that populations from drier environments had a greater probability of severe wilting under drought stress. Together, these patterns are consistent with a drought escape strategy rather than drought avoidance or drought tolerance.

**Conclusion:** Our results suggest that increasing aridity can favor integrated drought escape strategies characterized by rapid growth and early reproduction. These coordinated shifts reveal trade-offs between fast life-history strategies and investment in traits associated with drought resistance, highlighting how environmental gradients shape adaptive trait syndromes in wild plant populations.

## INTRODUCTION

Drought is a strong selective force shaping plant evolution, influencing trait diversification, life-history strategies, and species distributions across terrestrial ecosystems (Bray, 1997; Rauschkolb et al., 2022a; Stebbins, 1952). Climate change is expected to intensify drought regimes through increasingly erratic precipitation and elevated evaporative demand (Dai, 2013; IPCC, 2023). Understanding how plants respond to intensifying drought is critical for predicting evolutionary trajectories, informing conservation strategies, and identifying adaptive traits with potential applications in crop improvement.

Plants have evolved multiple strategies for coping with drought, broadly categorized as escape, avoidance, and tolerance (Figure 1; Fang & Xiong, 2015; Turner, 1986a). Drought escape is typical of annuals in environments where late-season drought terminates growth, and is achieved through rapid growth and early reproduction while soil moisture is still available (Franks, 2011; Rauschkolb et al., 2022b). Escape strategies are associated with traits such as faster growth, early maturity, elevated photosynthetic capacity, and reduced water-use efficiency (Blanco-Sánchez et al., 2022; Welles & Funk, 2021). In contrast, drought avoidance minimizes plant water stress through morphological and physiological adjustments that maintain tissue hydration, including increased root allocation and enhanced water-use efficiency (Farooq et al., 2009). Drought tolerance allows plants to function under low water potentials through mechanisms such as osmotic adjustment, osmoprotectant accumulation, and antioxidant defenses (Blum, 2010; Turner, 1986b). Because escape relies on rapid growth and high stomatal conductance, it often contrasts with the conservative water-use strategies characteristic of avoidance and tolerance. In addition, persistent selection for escape can relax selection on costly tolerance traits, reducing their prevalence in populations (Blumenthal et al., 2021; Haghpanah et al., 2024; Kooyers, 2015a).

**Figure 1:**
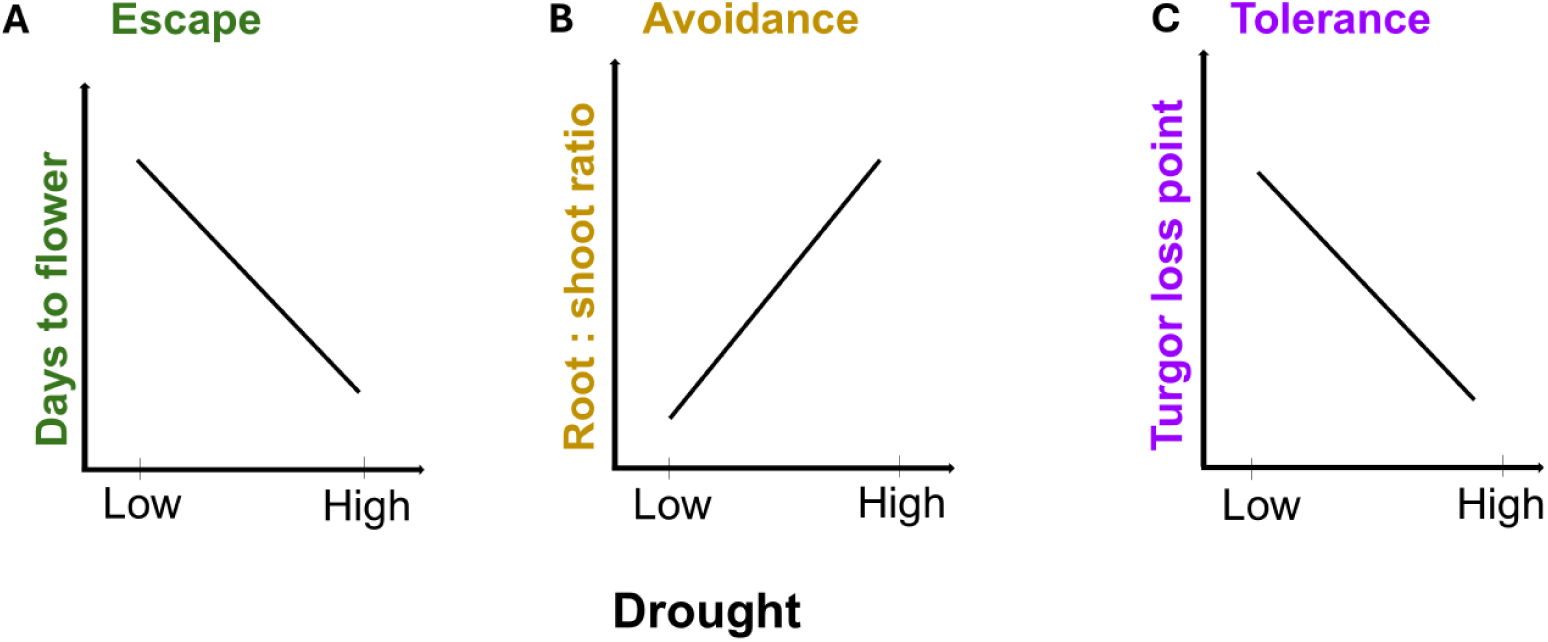
Conceptual representation of plant drought strategies along a drought gradient. Increasing drought stress is expected to promote earlier maturity in populations exhibiting an escape strategy (A), enabling completion of the life cycle before the onset of severe water limitation. In an avoidance response, plants are predicted to allocate more biomass to roots than to shoots (B), thereby enhancing water acquisition and reducing exposure to soil moisture deficits. The turgor loss point is a key proxy for tolerance, which is expected to be more negative under drought conditions (C), because it enables the maintenance of cellular turgor and physiological function at lower tissue water potentials.

Many studies focus on individual drought-associated traits, making it difficult to determine whether these traits evolve independently or as integrated strategies shaped by natural selection (Laughlin & Messier, 2015; Pivovaroff et al., 2016; Welles & Funk, 2021). Consequently, empirical evidence for coordinated variation among multiple drought-related traits across natural drought gradients remains relatively limited, particularly in wild annual plants (Burnette & Eckhart, 2021). In addition, accurately quantifying drought requires identifying environmental gradients that reflect the long-term cumulative water stress experienced by populations (Stephenson, 1998). Climatic moisture deficit (CMD), defined as the sum of the monthly difference between reference evaporative demand and precipitation, reflects long-term water limitation, with higher values indicating greater water deficit (Girvetz & Zganjar, 2014; Hargreaves, 1975). As CMD likely captures the long-term water limitations, variation in CMD across environments is expected to influence patterns of phenotypic change and determine which drought-adaptation strategies are favored. Evaluating suites of plant traits across CMD gradients can therefore provide insight into whether populations adopt strategies such as drought escape, avoidance, or tolerance through coordinated trait shifts, or whether traits respond independently.

Wild sunflower (*Helianthus annuus* L.) provides an excellent system for understanding the evolutionary basis of drought adaptation (Kantar et al., 2015; Marek, 2019; Seiler et al., 2023). Native to the semi-arid and arid regions of North America, wild *H. annuus* populations frequently colonize marginal habitats and experience substantial variation in moisture availability, seasonality, and evaporative demand (Heiser, 1951). Prior studies have documented extensive population-level differences in plant morphology, physiology, flowering phenology, and resource allocation patterns (Mandel et al., 2011; Putt, 1997; Todesco et al., 2020). This phenotypic diversity provides an excellent opportunity to examine how drought-related traits vary among populations and whether such variation aligns with distinct adaptive strategies across environmental gradients.

We used a common-garden experiment to study the evolutionary relationship between climatic moisture deficit and variation in drought-related traits across an aridity gradient in wild *H. annuus*. Specifically, we asked: (1) Does CMD represent a primary environmental axis structuring phenotypic variation among populations? (2) Which drought-adaptation strategies predominate across populations along the CMD gradient? and (3) Do growth, reproductive, and water-use traits exhibit trade-offs consistent with functional constraints associated with drought adaptation?

## METHODS

### Study System

Wild *Helianthus annuus* is a diploid (2n = 34), annual species native to North America and distributed across a broad range of environments from southern Canada to northern Mexico (Heiser et al., 1969; Seiler & Rieseberg, 1997). It is the wild progenitor of the domesticated sunflower (*H. annuus* L. var. macrocarpus), from which it remains genetically and ecologically distinct despite ongoing gene flow in some regions (Blackman et al., 2011; Harter et al., 2004). Unlike their domesticated counterparts, wild sunflowers are typically highly branched and produce smaller flower heads and seed (achenes) sizes (Seiler, 1992). Wild *H. annuus* also exhibits indeterminate flowering, extending several months into its growing season, typically from early spring to late summer across much of its native range (Presotto et al., 2011; Smith, 1978). As obligate outcrossers with a strong self-incompatibility system, wild sunflowers maintain high levels of genetic diversity within and among populations (Putt, 1997).

### Population Sampling

Between September and December 2022, we sampled 19 wild sunflower populations across Southern California (Figure 2, Appendix S1). To maximize genetic diversity, we sampled populations that had more than 30 mature plants and were located at least 3.2 km apart. From each population, we collected mature seed heads from 30 randomly selected mother plants, bagging each maternal line separately to preserve family structure. At each collection location, we recorded GPS coordinates using the GPS Camera 55 Field Survey mobile application (v 4.4.1). We used the ClimateNA web application to obtain high-resolution climate data (1 km) for Hargreaves climatic moisture deficit (mm) over the 30-year historical period (1990–2020) (Wang et al., 2016).

**Figure 2:**
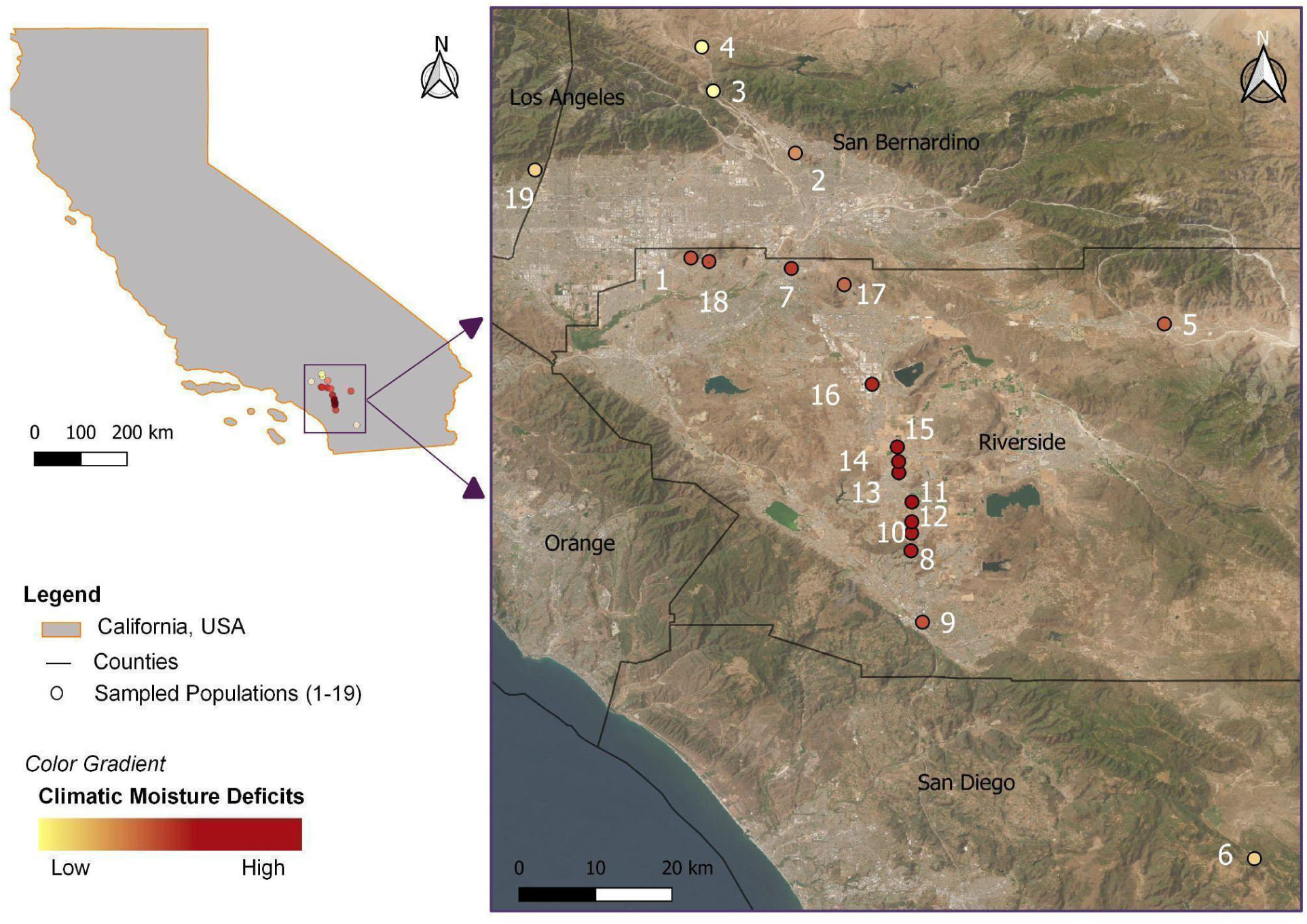
Distribution of sampled populations across Southern California. Sites (1–19; Appendix S1) are color-coded by Climatic Moisture Deficits from low (yellow) to high (red).

To characterize local edaphic conditions as a second potential driver of adaptive variation, we collected five soil samples per population at 10–25 cm depth across a linear 150 ft transect. Soil samples were oven-dried at 70 °C for 48 hours, sifted through a 4000-micron sieve, pooled by population, and sent to Denele Analytical Inc. for comprehensive soil analyses, including soil pH, electrical conductivity, nitrate nitrogen, phosphorus, boron, sulfate, micronutrients (Zn, Fe, Cu, Mn), organic matter, carbonates, cation exchange capacity, nitrogen holding capacity, exchangeable sodium percentage, and sodium adsorption ratio.

### Plant Establishment and Common Garden

From each population, we randomly selected 12 maternal lines to be included in the common garden. From each maternal line, five seeds were mechanically scarified by cutting the blunt end of the achene to breach the pericarp opposite the embryo, enabling rapid imbibition and breaking physical dormancy. We germinated scarified seeds on filter paper in a petri dish at room temperature (∼21 °C) and in the dark. After 48 hours, we changed the filter paper and removed seed coats to reduce microbial growth and enhance germination. The petri dishes were then kept in the dark for another 72 hours, followed by 48 hours of light exposure. One seedling per germinated maternal line was transplanted into cell insert packs (5.02 cubic inches volume), containing Pro-Mix Biofungicide and Mycorrhizae (75-85% sphagnum peat base and introductory fertilizer). The seedlings were grown in a light cart (1,900-lumen tube lights) at room temperature for three weeks, with trays regularly shuffled to minimize positional effects, before being transplanted into 15 cm standard pots (1835 cm^3^) containing UC soil mix III.

Plants were grown in a lath house at the University of California, Riverside, and irrigated daily through drip lines until saturation for the first four weeks of transplanting. Thereafter, irrigation was reduced to twice per week until the drought phase, during which drip lines were removed to impose water stress. Plants were fertigated twice during the experiment using a 60 g/L fertilizer solution (N:P:K = 21:5:20) at a 0.01% application rate. Plant pots were randomly shuffled across the two bench spaces multiple times throughout the growth period to minimize the positional effect.

### Phenotypic Measurements

During the common garden experiment, we measured pre-drought and post-drought phenotypes. Among the pre-drought phenotypes, we measured plant height, stem diameter, leaf number, specific leaf area, fresh, turgid, and dry weight of leaves, relative water content of leaves, number of branches, buds, and flowers, days to first bud and flower, petiole length and diameter, leaf angle, leaf temperature, and leaf chlorophyll content. Plant height was recorded from the soil surface to the highest peduncle due to the lack of apical dominance in wild sunflowers at 64 and 121 days after germination (hereafter referred to simply as days). Stem diameter was recorded 2 cm above the soil using digital vernier calipers at 64, 99, and 121 days. We counted leaf numbers, excluding small sessile leaves from the main stem. Similarly, we selected a healthy and recently expanded leaf at 100 days after germination (referred to simply as days from onwards), and calculated its area using ImageJ. We then recorded water-saturated fresh weight and oven-dry weight (60°C for 24 hours) to calculate SLA and LDMC using the following equations.

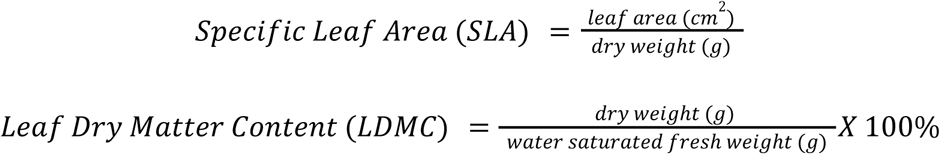

To measure relative water content (RWC), we collected 2.5 cm diameter leaf discs of a mature leaf at 156 days. Fresh weight was recorded before leaves were hydrated for 24 hours to obtain turgid weight, and then dried at 60°C for 24 hours to determine dry weight.

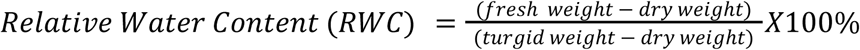

For phenological traits, we noted days to first budding and flowering, and when more than two-thirds of the plants had flowered (148 days), we recorded the number of buds, flower heads, and the final flowering status. At this stage, we counted branch numbers and measured the length and diameter of a randomly selected healthy leaf petiole. At 142 days, we randomly selected a healthy mature leaf to measure photosynthetic phenotypes, including chlorophyll concentration, leaf temperature, leaf thickness, and leaf angle using the MultiSpeQ v2.0 (Jamil et al., 2026; Kuhlgert et al., 2016).

At 103 days, leaf water potential at turgor loss (π_tlp_) was characterized as cellular osmotic potential (Bartlett et al. 2012) for a subset of the plants (three randomly selected individuals per population). Leaf discs (6.2 mm diameter) were excised from leaves, wrapped in aluminum foil, and submerged in liquid N_2_ for 2 min to rupture the cellular structure. Upon removal from liquid N_2_, leaf discs were punctured ten times with sharp-tipped forceps before sealing in the osmometer chamber (Model 5600; Wescor Inc, Logan, Utah, USA), for measurement of osmolality (mOsm kg^-1^). Final osmometer readings were converted to π_tlp_ using the van’t Hoff equation (Bartlett et al., 2012; Maréchaux et al., 2015).

To impose water stress, drip irrigation was discontinued at 161 days, and leaf water potential and wilting score were measured 7 days later as post-drought phenotypes. To measure leaf water potential, we clipped one mature leaf per plant in the pre-dawn. We kept them sealed in plastic bags and stored them inside a cooler to minimize water loss during transport to the lab, where their leaf water potential was measured using a pressure chamber (Model 1000; PMS Instruments, Albany, Oregon, USA) (Rodriguez-Dominguez et al., 2022). On the same day, we scored the plants for wilting on a scale of 1 (minimal wilting) to 5 (severe wilting).

### Statistical Analysis

All statistical analyses were conducted in R v4.4.3 (R Core Team, 2000). To characterize edaphic variation, soil variables were standardized before analysis and summarized using principal component analysis (PCA) to reduce dimensionality and capture major axes of soil variation. Patterns of covariation among plant traits and environmental variables were visualized using pairwise plots generated with the GGally package (Schloerke et al., 2021).

To understand the role of environmental variables shaping trait variation, we fitted a series of models including CMD and soil variation summarized by soil PCs (PC1 and PC2). Model performance was evaluated using Akaike Information Criterion (AIC) and likelihood ratio tests, where models with only CMD were consistently the best fit (Appendix S2), especially after correcting for multiple comparisons using a Bonferroni correction for 20 tests and adjusting our alpha value to 0.0025. Furthermore, we used variance partitioning with the varpart function in the vegan package (Oksanen et al., 2009) to estimate the variation in traits explained by CMD and the two soil PCs (Appendix S3).

We modeled trait responses using regression frameworks that depended on the distribution of each response variable. Continuous traits were analyzed using linear mixed-effects models assuming Gaussian error distributions. Binary outcomes (flowering probability) were modeled using binomial logistic regression, while ordinal traits (wilting score) were analyzed using cumulative link models. Traits with count numbers were modeled using generalized linear mixed-effects models with either Poisson (number of flowers) or negative binomial (number of buds) error distributions, depending on the degree of overdispersion.

All mixed-effects models were fitted using the *lme4* package (Bates et al., 2015), with one or more environmental variables as fixed effects and population origin as a random effect (e.g., Trait ∼ CMD + [1|Population ID]). For the analysis of wilting probability, CMD was standardized before modeling by subtracting the mean and dividing by the standard deviation to ensure comparable effect sizes across models. Predicted probabilities for each wilting score were generated across the CMD gradient, and 95% confidence intervals were obtained by simulating model coefficients from the variance–covariance matrix of the fitted models.

Due to the asymmetry in the distances between populations in our collections, where populations are closer together in drier regions, we grouped geographically proximate populations (IDs 8, 10, and 12 as one group, and 13, 14, and 15 as another; Figure 2, Appendix S4) to explore potential bias arising from pseudoreplication. We then fitted a series of models using these grouped populations to assess the robustness of our results. Model outcomes were qualitatively consistent across grouping schemes, indicating that the observed patterns were not driven by sampling imbalance (Appendix S4).

Because our preliminary analyses showed strong correlations between plant size and other traits, we wanted to know whether these trait correlations vary across populations with different CMD. To test for this, we compared additive and interaction models with the predictors initial plant height and CMD using likelihood ratio tests. We found limited evidence for significant interactions, indicating that the effect of initial plant height on other traits was mostly consistent across our drought gradient (Appendix S5).

## RESULTS

### Soil properties

A PCA based on 20 soil variables identified two major axes of variation (Appendix S6). Soil PC1 (24.6% of variance) represented a broad gradient in soil chemical composition and nutrient status. Positive scores on this axis corresponded to soils with higher nutrient-holding capacity, cation exchange capacity, and base cations (calcium, magnesium), while negative scores reflected soils with relatively higher micronutrient availability (iron, copper, boron). Soil PC1 was positively associated with chlorophyll content (β = 0.337), indicating higher chlorophyll in populations originating from more nitrogen-retentive soils, consistent with known relationships between nitrogen availability and leaf chlorophyll content (Evans, 1989). Soil PC2 (19.9% of variance) captured variation in soil salinity associated with sodium accumulation, driven by several variables (exchangeable sodium percentage, sodium adsorption ratio, and sodium cations).

### Traits and environmental covariation

Pairwise correlations revealed extensive covariation among plant traits and environmental variables (Figure 3, Appendix S7). Plant height was positively correlated with SLA, days to first flowering, and wilting score and negatively correlated with root: shoot ratio, indicating a shared axis of trait variation. Similarly, CMD was strongly associated with multiple plant traits. Although not associated with soil PC2 (r = 0.35, P > 0.05), CMD was strongly associated with soil PC1 (r = 0.35, P < 0.01), which is consistent with increasing base cation accumulation under greater aridity (Appendix S8). Despite this relationship, incorporating soil variables into models of trait variation did not consistently improve fits relative to models with CMD alone (Appendix S2), and variance partitioning indicated that CMD explained a larger portion of trait variation than soil PCs across most phenotypes (Appendix S3). We therefore present the results of trait analyses that used CMD as the only environmental predictor in the sections below.

**Figure 3:**
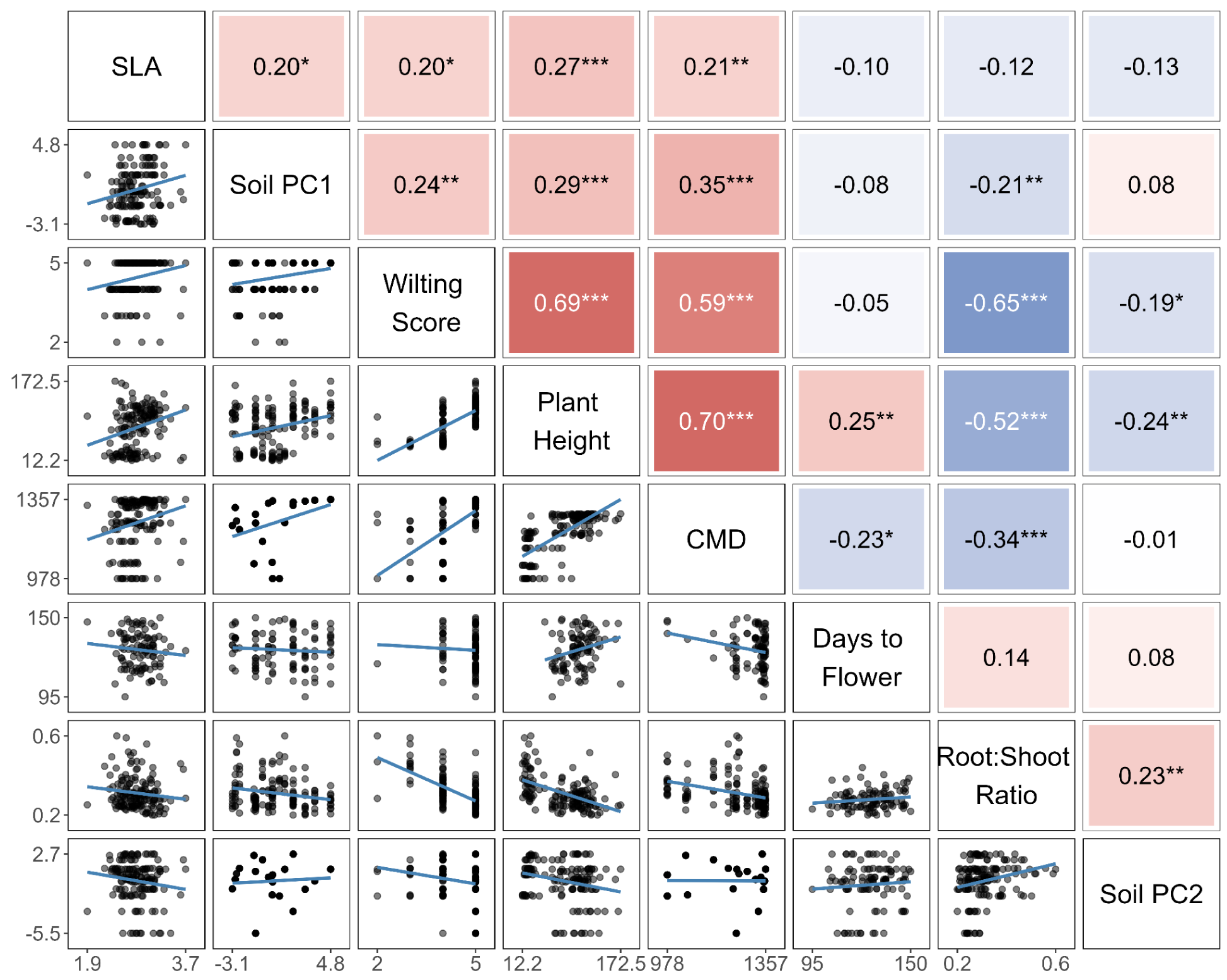
Pairwise relationships among climatic moisture deficit (CMD), soil PC1, Soil PC2, and plant traits (height at 121 days, specific leaf area [SLA], days to flowering, root: shoot ratio, and wilting score). Upper panels show Spearman correlation coefficients, while lower panels show bivariate scatterplots with fitted lines. Colors indicate the direction of the correlation (red as positive and blue as negative), and color intensity reflects the strength of the correlation. Asterisks indicate significance level: *** (*P*< 0.001), ** (*P*< 0.01), * (*P*< 0.05). Traits are clustered based on their correlation coefficient (r). CMD is associated with a coordinated shift toward taller plants with higher SLA, earlier flowering, reduced root allocation, and increased wilting. Furthermore, CMD is significantly correlated with Soil PC1. See Appendix S8 for pairwise correlations for all the traits.

### Coordinated shifts in growth and reproductive strategy across the aridity gradient

Regression analyses showed substantial variation in plant growth and reproductive traits across the aridity gradient. Populations originating from higher CMD environments were taller (R^2^= 0.465, P < 0.001; Figure 4a) and exhibited faster height growth rates (R^2^ = 0.233, P < 0.001; Figure 4b) in the common garden. Specific leaf area increased modestly with CMD (R^2^ = 0.042, P = 0.011; Figure 4c), and leaf angle became more horizontal (R^2^ = 0.107, P = 0.005; Figure 4d). Leaf temperature also increased along the gradient (R^2^ = 0.040, P = 0.035; Appendix S9b), although observed values likely remained within the thermal range associated with active photosynthesis (Yamori et al., 2014).

**Figure 4:**
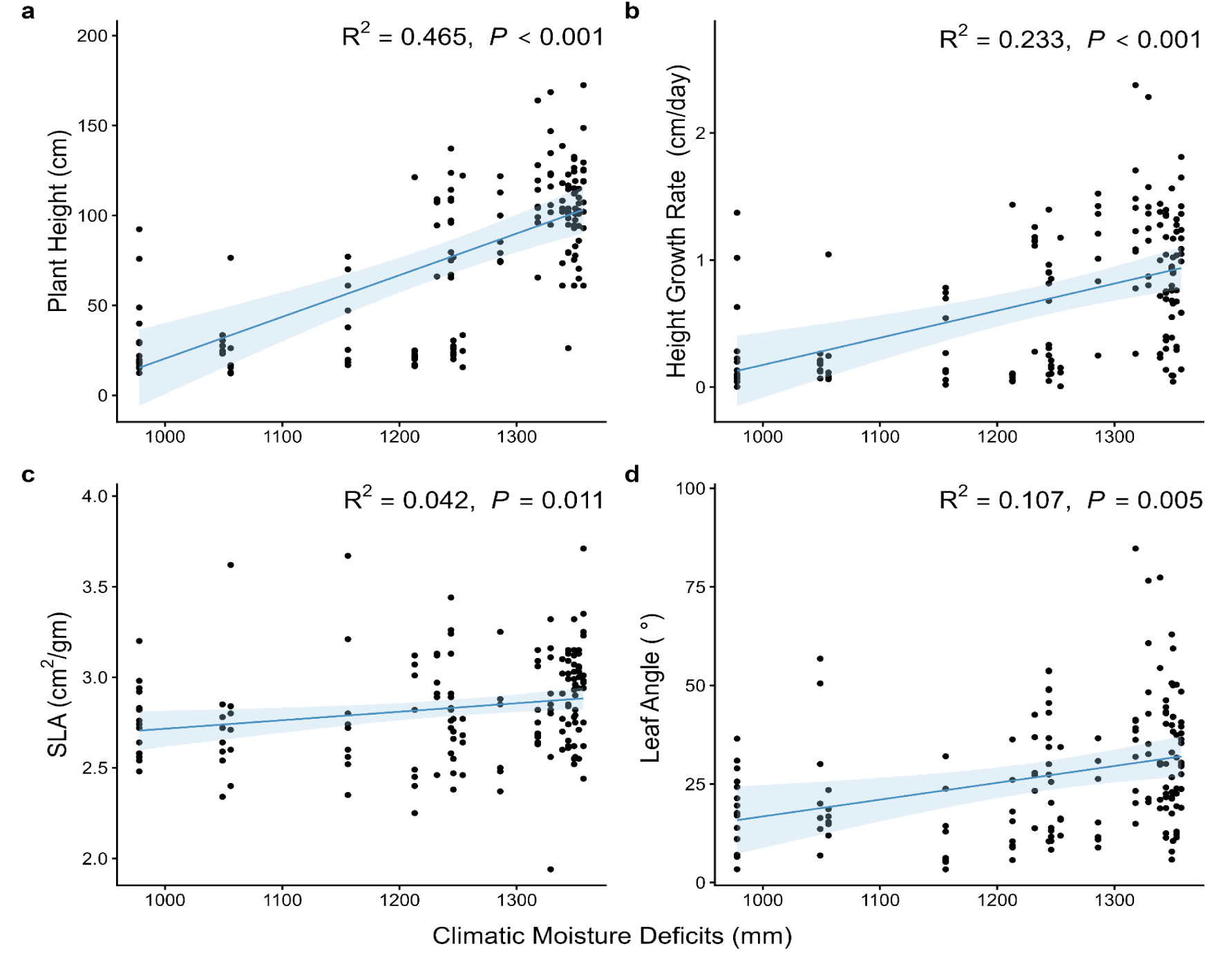
Relationships between climatic moisture deficit (CMD) at population origin and plant growth traits measured in a common garden. (a) Final plant height (cm), (b) height growth rate (cm day⁻¹) between 64 and 149 days after germination, (c) specific leaf area (SLA; cm² g⁻¹), and (d) leaf angle (°). Lines represent fitted mixed-effects model predictions with 95% confidence intervals. Points represent individual plants. Marginal R² and P-values for the fixed effect of CMD are shown in each panel. Across the CMD gradient, populations from drier origins grew taller, faster, and developed broader, more horizontally oriented leaves.

Reproductive traits showed similarly pronounced shifts. Days to first flowering declined with increasing CMD (R^2^ = 0.068, P = 0.019; Figure 5a), indicating earlier flowering in populations from drier origins. Along the increasing aridity gradient, the probability of flowering increased strongly (R^2^ = 0.339, P < 0.001; Figure 5b), as did the reproductive output, including the number of buds (R^2^ = 0.636, P = 0.006; Figure 5c) and flower heads (R^2^ = 0.549, P < 0.001; Figure 5d). Together, these patterns indicate coordinated increases in plant stature, growth rate, flowering likelihood, and reproductive output among populations from greater moisture deficit environments.

**Figure 5:**
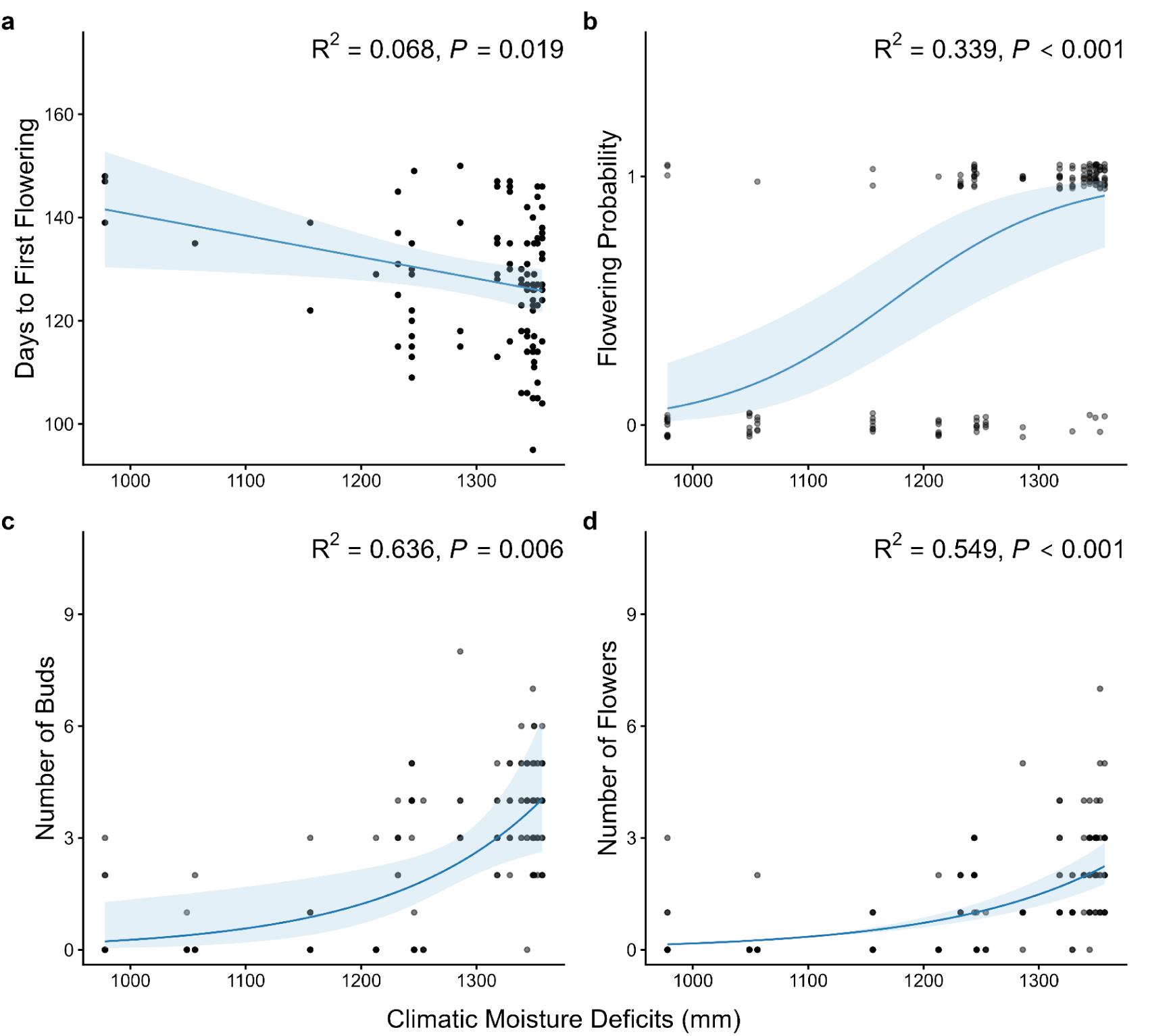
Relationships between climatic moisture deficit (CMD) at population origin and reproductive phenology and output measured in a common garden: (a) days to first flowering, (b) probability of flowering, (c) number of buds, and (d) number of flowers. Lines represent fitted mixed-effects model predictions with 95% confidence intervals, and points represent individual plants. Days to first flowering was modeled using linear mixed-effects models, whereas flowering probability (binomial), number of buds (negative binomial), and number of flowers (Poisson) were modeled using generalized linear mixed-effects models. Marginal R² and P-values for the fixed effect of CMD are shown in each panel. Across the CMD gradient, populations from drier origins flowered earlier and invested more heavily in reproduction.

### Tradeoff in resource allocations and water status

The rapid growth and early maturity among populations from high CMD environments were accompanied by coordinated shifts in plant structure and biomass allocation (Figure 6, Figure S3, Appendix S8). Leaf and structural investment declined with increasing CMD, with thinner leaves (R² = 0.246, P = <0.001; Appendix S9a), lower leaf dry matter content (R² = 0.042, P = 0.01; Figure 6b), alongside reduction in stem diameter (R^2^ = 0.302, P < 0.001; Appendix S10a), petiole length (R^2^ = 0.267, P < 0.001; Appendix S10c), and petiole cross-sectional area (R² = 0.281, P < 0.001; Appendix S10d). Allocation to belowground tissues also declined with increasing CMD, with lower root: shoot biomass ratios in populations from drier origins (R² = 0.112, P = 0.032; Figure 6a).

**Figure 6:**
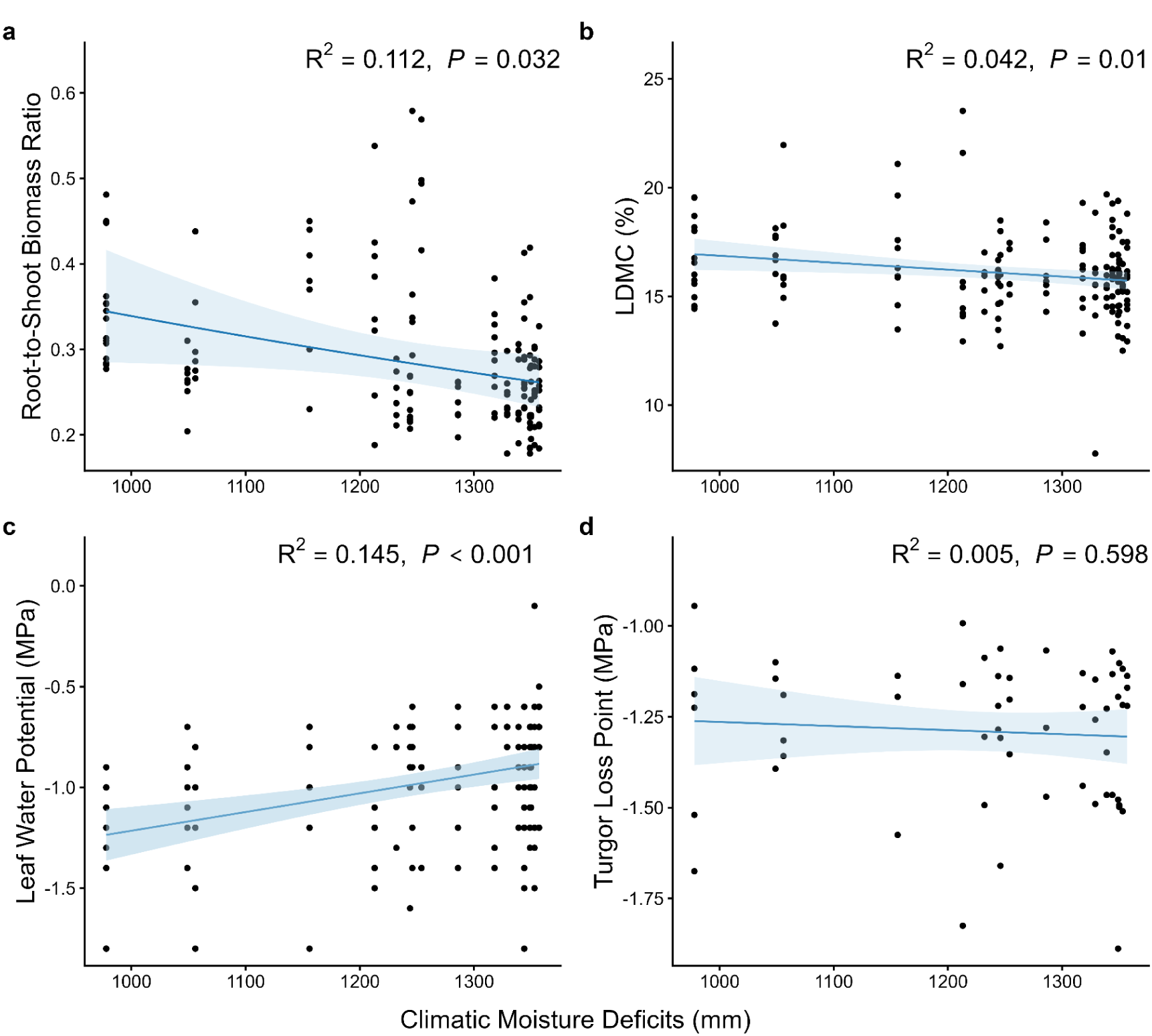
Relationships between climatic moisture deficit (CMD) at population origin and drought-related functional traits measured in a common garden: (a) root-to-shoot biomass ratio, (b) leaf dry matter content (LDMC; %), (c) turgor loss point (MPa), and (d) leaf water potential (MPa). Lines represent fitted mixed-effects model predictions with 95% confidence intervals, and points represent individual plants. Marginal R² and P-values for the fixed effect of CMD are shown in each panel. Across the CMD gradient, populations from drier origins exhibited lower root allocation and reduced LDMC, maintained higher leaf water potentials, and showed little variation in turgor loss point (measured on a subset of individuals).

Patterns in plant water status were broadly consistent with reduced drought tolerance. Ordinal logistic models predicted that wilting severity increases along the CMD gradient (Figure 7), with the probability of the most severe wilting category (score 5) increasing toward higher CMD origins, whereas low-wilting categories (scores 2–3) remained uncommon across the gradient. Consistent with this pattern, relative water content declined slightly with increasing CMD (R² = 0.042, P = 0.051; Figure S3d), and plants from drier origins maintained higher (less negative) leaf water potentials (R² = 0.145, P < 0.001; Figure 6c). In contrast, turgor loss point showed no association with CMD (R² = 0.005, P = 0.598; Figure 6d) when measured at 103 days in a subset of individuals (n = 57).

**Figure 7:**
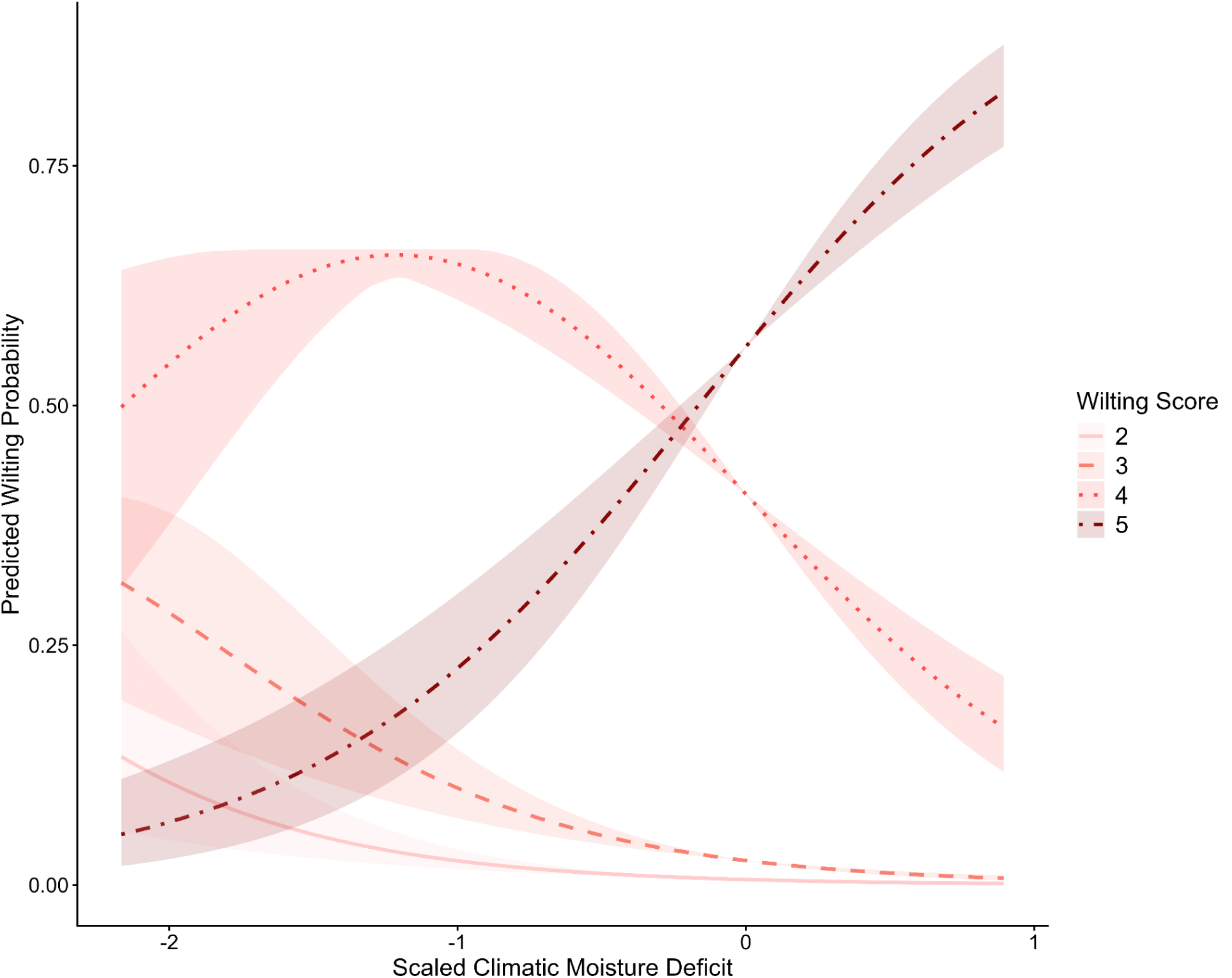
Predicted probabilities of wilting score categories across the climatic moisture deficit (CMD) gradient. Lines show fitted values from an ordinal logistic regression including standardized CMD as a predictor. Shaded bands indicate 95% confidence intervals. Line types and colors denote wilting categories (2 = least wilted; 5 = most severely wilted). The probability of severe wilting increased with CMD.

## DISCUSSION

### Climatic moisture deficit as a major environmental gradient shaping trait variation

Using a common-garden experiment across populations of wild *Helianthus annuus*, we examined how climatic moisture deficit (CMD) relates to phenotypic variation in drought-related traits across an aridity gradient. Our results are consistent with CMD acting as an important environmental gradient associated with coordinated shifts in plant growth, phenology, and resource allocation. Several traits, including plant height, growth rate, flowering phenology, leaf morphology, and biomass allocation, varied systematically with CMD, indicating that multiple aspects of plant function change along the aridity gradient. These coordinated shifts reflect changes across a suite of traits rather than a single trait in isolation (Díaz et al., 2016; Reich, 2014)

Although CMD explained variation in several phenotypes, substantial population-level variation remained. This pattern is consistent with the high genetic diversity typical of wild sunflower populations (Mandel et al., 2011; Park & Burke, 2020) and suggests that responses to aridity likely reflect both broad climatic gradients and local selective histories. While soil properties also varied along the CMD gradient, variance partitioning and model comparisons indicate that they explained smaller and less consistent proportions of trait variation than CMD. These results support CMD as a key environmental parameter underlying phenotypic differentiation among populations, while highlighting the contribution of additional sources of variation along the CMD gradient.

### Drought escape and life-history strategies

Multiple phenotypes point toward drought escape as a primary adaptive strategy among populations originating from arid environments. Although drought adaptation is often discussed in terms of resistance or tolerance mechanisms, escape strategies are frequently favored by arid annual plants experiencing terminal seasonal drought (Cohen, 1976; Franks, 2011). In our study, plants originating from drier regions flowered earlier and were more likely to reach reproductive maturity during the growing season. Earlier flowering allows plants to complete reproduction before severe water limitation develops later in the season and is widely recognized as a key component of drought escape in annual species (Blanco-Sánchez et al., 2022; Monroe et al., 2018; Shavrukov et al., 2017).

We also found that plants originating from more arid sites attained greater height, and taller plants were more likely to flower. Furthermore, those plants have exhibited higher SLA, more horizontal leaf angles, and slightly elevated leaf temperatures, traits commonly associated with rapid resource acquisition and fast early-season growth. Together, these patterns suggest a life-history strategy emphasizing rapid early growth and accelerated reproduction, enabling plants to complete their life cycle before water limitation intensifies.

### Trade-off in growth and drought vulnerability

While populations from drier environments exhibited traits consistent with drought escape, these shifts were accompanied by reduced investment in traits associated with structural support and drought buffering. Populations originating from drier environments generally exhibited lower root-to-shoot biomass ratios, reduced relative water content, and more severe wilting at less negative leaf water potentials, indicating a reduced capacity to maintain water status as soil moisture declines. Such patterns are consistent with life-history strategies that prioritize rapid growth and early reproduction over investment in long-term physiological resilience (Reich, 2014; Wu et al., 2010).

Despite strong associations between CMD and many drought-related traits, we did not detect a significant relationship between CMD and turgor loss point in a subset of individuals for which this trait was measured. Turgor loss point is widely considered a key physiological indicator of drought tolerance (Bartlett et al., 2014; Maréchaux et al., 2015), and the absence of a CMD relationship suggests that costly physiological tolerance mechanisms may be less strongly favored when drought escape strategies dominate. Similar patterns have been documented in other annual systems, where earlier flowering and accelerated growth reduce the selective advantage of investment in drought tolerance traits (Burnette & Eckhart, 2021; Franks, 2011; Geber & Dawson, 1990; Kooyers, 2015b).

Collectively, these results suggest that adaptation along the aridity gradient involves coordinated shifts in life-history, structural, and water-use traits. In this system, selection for rapid growth and early reproduction appears to constrain investment in traits associated with drought avoidance or tolerance, producing integrated trait strategies that reflect trade-offs between growth and drought resilience. Comparable coordination among drought escape, avoidance, and tolerance strategies has been documented across Mediterranean annuals, desert annuals, and grassland species, where increasing aridity often favors earlier flowering, faster growth, and reduced investment in structural or water-conserving traits (Blanco-Sánchez et al., 2022; Franks et al., 2007; Hamann et al., 2018; Welles & Funk, 2021).

### Implications for evolution and crop improvement

Our findings suggest that wild *H. annuus* populations from drier sites have evolved a coordinated suite of traits consistent with a drought escape–dominated life-history strategy, combining rapid growth, early flowering, and reduced investment in structural tissues and dehydration tolerance. This pattern highlights how aridity can shape integrated whole-plant strategies rather than isolated traits.

From an applied perspective, these naturally evolved trait combinations represent a valuable reservoir of adaptive variation for sunflower improvement. Traits such as early flowering, rapid growth, and high SLA observed in arid-adapted populations could be leveraged to enhance performance in dryland or rainfed systems by allowing crops to complete reproduction before severe water limitation occurs. Such escape-based strategies provide a complementary alternative to classical drought tolerance approaches, which can sometimes incur yield penalties due to high metabolic or structural costs (Blanco-Sánchez et al., 2022; Down-to-earth, 2024; Shavrukov et al., 2017). At the same time, the trade-offs documented here highlight the importance of considering coordinated trait syndromes when incorporating wild variation into crop improvement programs. Integrating traits that influence growth, phenology, and water use will be essential for balancing productivity and stress resilience under increasingly variable climatic conditions (Soriano et al., 2004). More broadly, these results illustrate how studying natural adaptation across environmental gradients can inform both our understanding of plant evolutionary responses to aridity and the development of crop varieties better suited to future climates.

## CONCLUSIONS

Our results show that climatic moisture deficit is strongly associated with coordinated variation in growth, phenology, and resource allocation among populations of wild sunflowers. Populations from more arid environments exhibited traits consistent with a drought escape strategy, including earlier flowering, rapid growth, and traits linked to fast resource acquisition. These shifts were accompanied by reduced investment in traits associated with drought buffering, highlighting trade-offs between rapid life-history strategies and physiological drought tolerance. Together, these findings suggest that aridity can shape integrated plant strategies rather than individual traits in isolation. Broadly, these results highlight the value of studying natural variation across aridity gradients for understanding plant adaptation and for identifying traits that may improve crop performance under increasing drought stress.

## ACKNOWLEDGMENTS

We thank Dr. Alan Brelsford for his valuable guidance throughout this work. We are also grateful to Bhawana Acharya, S. Gangothri, Teddy Reitman, Conner Lay, Brooke Wallasch, Aurora Cullen, Natthinee Sutjaitham, Alana Yamauchi, Mariana Moreno, Brinda Avadhanam, Narayan Bhusal, and Dr. Marko Spasojevic for their assistance and support across various stages of the project. This work was supported by the OASIS program through funding awarded to K.O.

## AUTHOR CONTRIBUTIONS

RN and KLO designed the study. RN, BRS, and KLO collected seeds. RN, BRS, LSS, and KLO phenotyped the common garden. RN and KLO analyzed the data. RN drafted the manuscript. All authors read, edited, and approved the final manuscript.

