## Supplementary Figures for "Drought escape as an adaptive strategy across an aridity gradient in wild sunflower"

### Appendix S6: Principal Component Analysis of soil variables

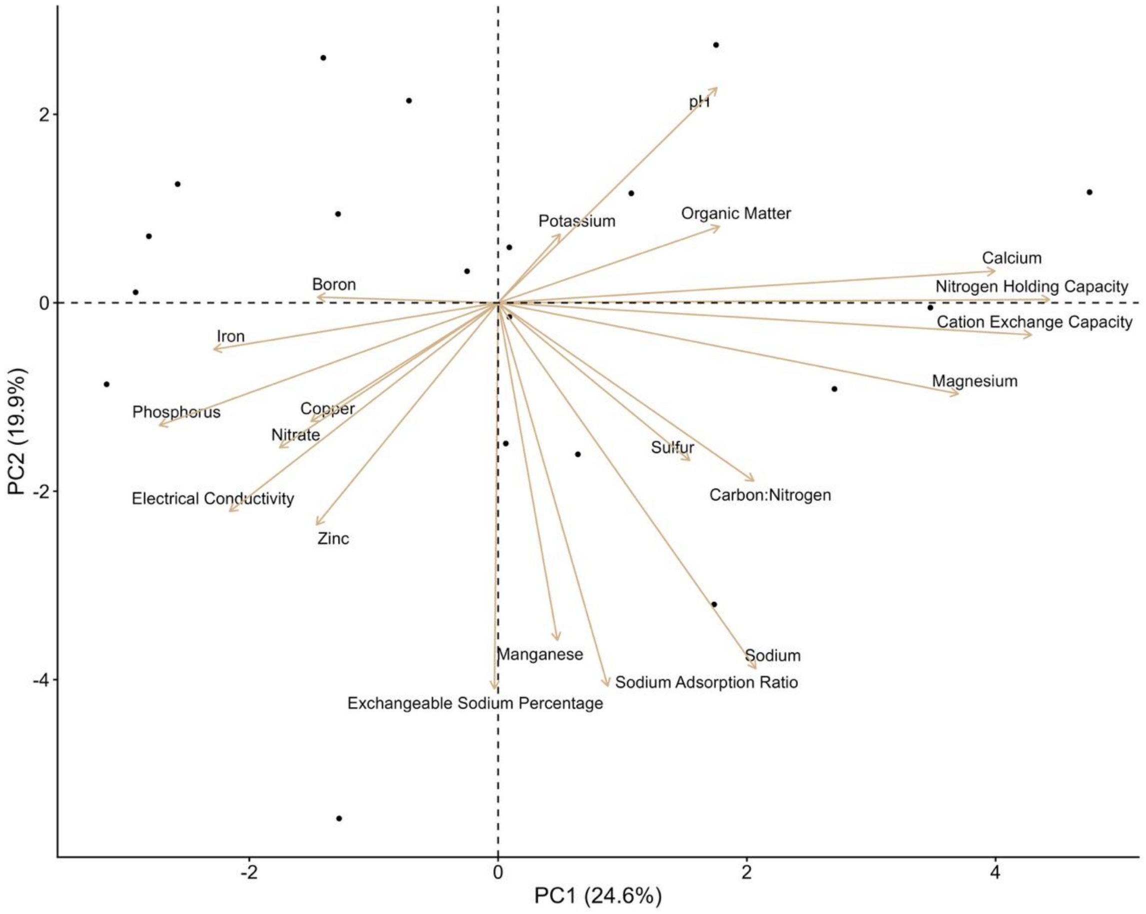


**Appendix S6** Principal component analysis (PCA) of soil properties across populations. Arrows indicate variable loadings on PC1 (24.6%) and PC2 (19.9%). PC1 primarily reflects a gradient from soils with higher nutrient retention and base cations to soils with higher micronutrients, whereas PC2 captures variation in soil pH and organic matter with sodium-related soil properties.

Appendix S7: Pairwise correlations among plant traits and climatic moisture deficit

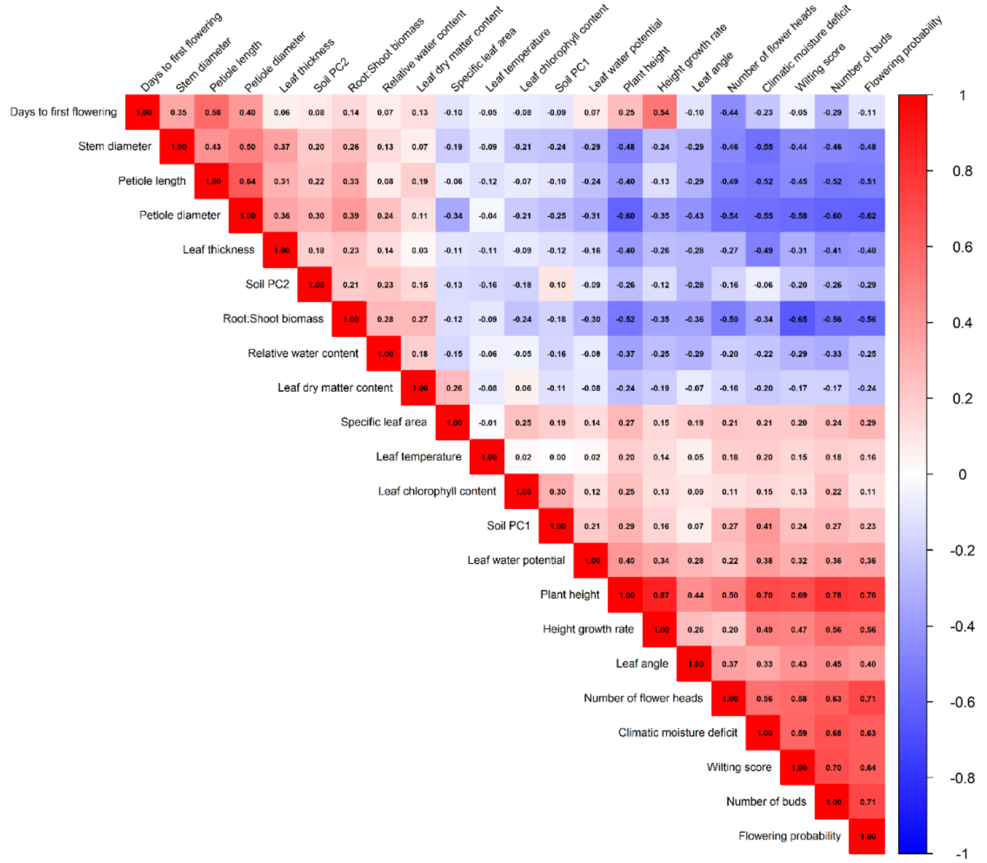


**Appendix S7:** Pairwise Pearson correlations among plant traits, climatic moisture deficit (CMD), and soil principal components (PC1 and PC2). The upper triangle shows correlation coefficients, with colors indicating the direction and strength of relationships (red = positive, blue = negative).

Appendix S8: Relationships between CMD and soil properties of population origins

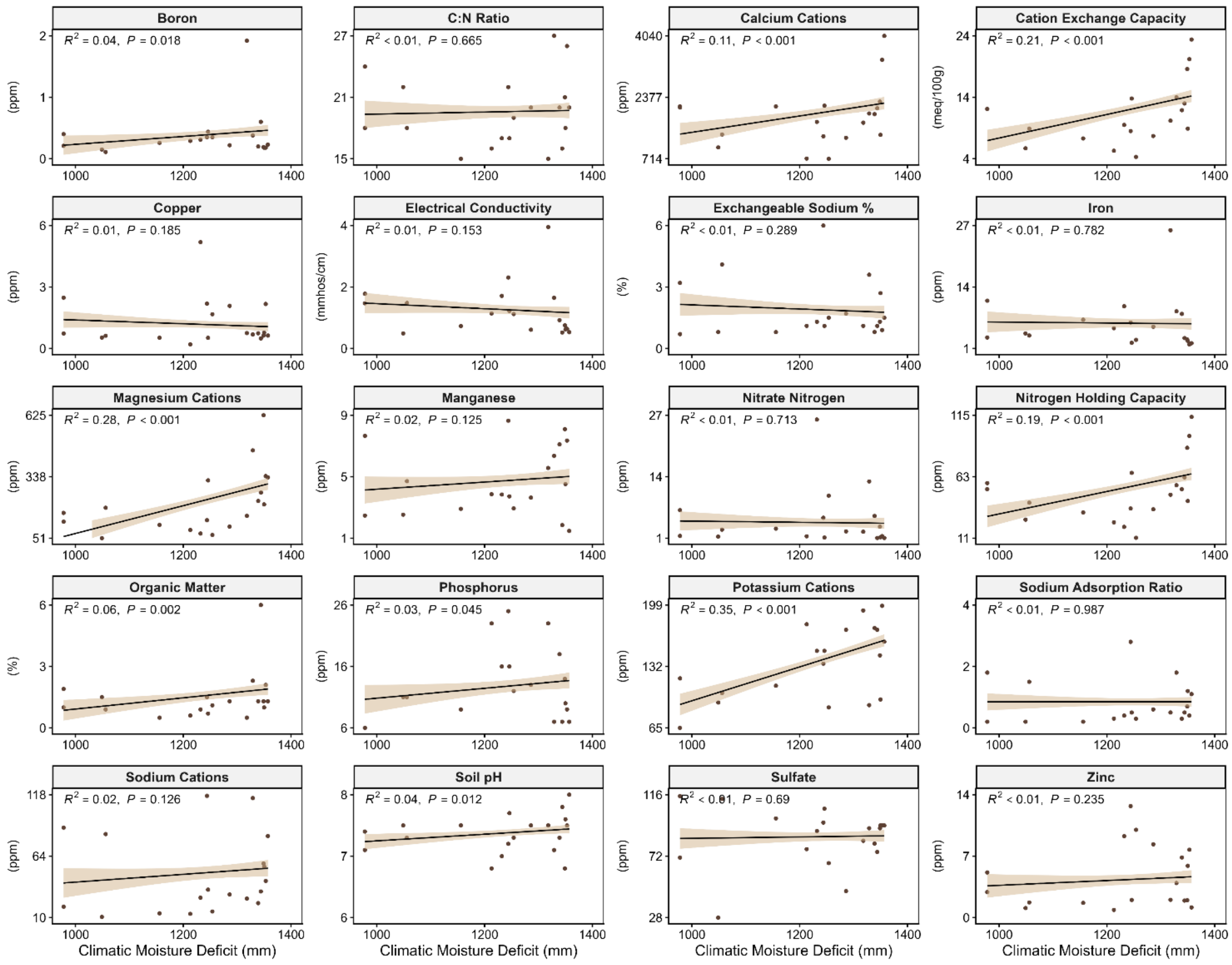


**Appendix S8:** Relationships between CMD and soil properties of population origins. Solid lines show fitted values from linear models with shaded areas indicating 95% confidence intervals. Higher CMD was strongly and significantly associated with increased potassium and magnesium cations, cation exchange capacity, and nitrogen holding capacity.

Appendix S9: Relationships between CMD at population origin and plant structural traits measured in a common garden.

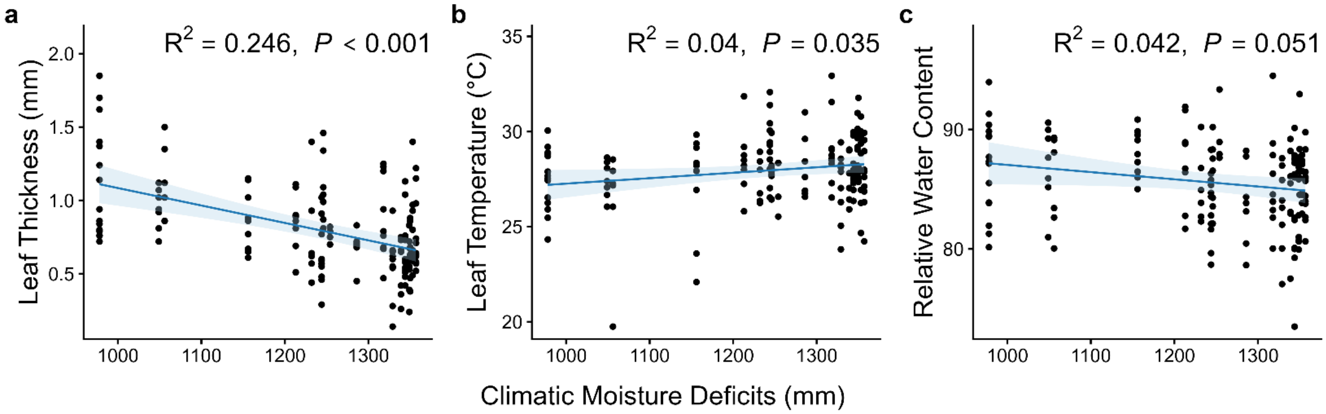


**Appendix S9:** Relationships between climatic moisture deficit (CMD) at population origin and leaf functional traits measured in a common garden: (a) leaf thickness (mm), (b) leaf temperature (°C), and (c) relative water content (%). Lines represent fitted mixed-effects model predictions with 95% confidence intervals, and points represent individual plants. Marginal $R^2$ and P-values for the fixed effect of CMD are shown in each panel. Across the CMD gradient, populations from drier origins had thinner leaves, slightly higher leaf temperatures, and modest declines in relative water content.

Appendix S10: Relationships between CMD at population origin and plant structural traits measured in a common garden.

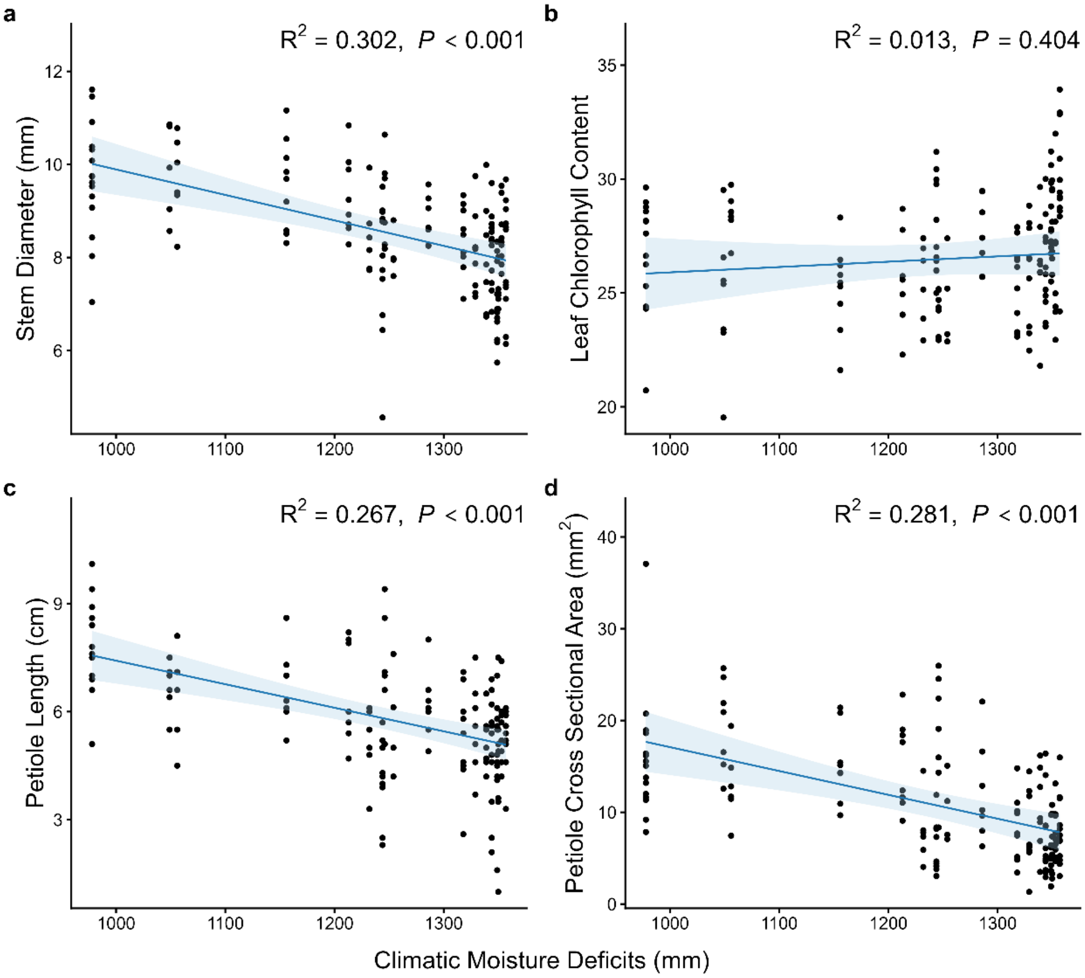


**Appendix S10:** Relationships between climatic moisture deficit (CMD) at population origin and plant growth and leaf structural traits measured in a common garden: (a) stem diameter (mm), (b) Leaf chlorophyll content, (c) petiole length (cm), and (d) petiole cross-sectional area ($mm^2$). Lines represent fitted mixed-effects model predictions with 95% confidence intervals, and points represent individual plants. Marginal $R^2$ and *P*-values for the fixed effect of CMD are shown in each panel. Across the CMD gradient, populations from drier origins exhibited smaller stem diameters, shorter petioles, and reduced petiole cross-sectional area, while chlorophyll content showed little relationship with CMD.
